# Radiation-induced rescue effect on human breast carcinoma cells is regulated by macrophages

**DOI:** 10.1101/2023.08.02.551610

**Authors:** Spoorthy Pathikonda, Li Tian, Shuk Han Cheng, Yun Wah Lam

## Abstract

The susceptibility of cancer cells to DNA damages is influenced by their microenvironment. For example, unirradiated neighbors of irradiated cells can produce signals that reduce DNA damages. This phenomenon, known as Radiation-Induced Rescue Effect (RIRE), has profound implications on the efficacy of radiotherapy. Using bystander cells cocultured with mock-irradiated cells as a control, we demonstrated, for the first time, two types of RIRE. Conditioned medium from naïve bystander cells, i.e., cells not exposed to irradiated cells, could mitigate UV-induced DNA damages in human breast carcinoma MCF7 cells, as judged by phospho-H2AX and 53BP1 immunostaining. This protective effect could be further enhanced by the prior treatment of bystander cells with factors from UV-irradiated cells. We named the former effect “basal RIRE” and the latter “active RIRE” which were cell type-dependent. As bystanders, MCF7 showed a significant active RIRE, whereas THP1-derived macrophages showed a strong basal RIRE but no active RIRE. Interestingly, RIRE of macrophages could further be modulated by polarisation. The basal RIRE of macrophages was abolished by M1 polarisation, while M2 and Tumour Associated Macrophages (TAM) demonstrated pronounced basal and active RIRE. When mixtures of MCF7 cells and polarised macrophages were used as bystanders, the overall RIRE was dictated by macrophage phenotypes: RIRE was suppressed by M1 macrophages but significantly enhanced by M2 and TAM. This study shows a previously unappreciated role of the innate immune system in RIRE. Depending on polarised phenotypes, macrophages in the tumour microenvironment can interfere with the effectiveness of radiotherapy by adjusting the RIRE magnitudes.

## Introduction

DNA damage response (DDR) is indispensable in maintaining genomic stability. It is activated when cells are exposed to DNA damaging agents such as ionizing or non-ionizing radiation and chemotherapeutic drugs (Li et al. 2021). Among these genotoxic challenges, ultraviolet light (UV) is one of the most prevalent, inducing up to 10^5^ DNA lesions in a human cell per day (Hoeijmakers 2009). The formation of UV-induced DNA adducts, predominately cyclobutane pyrimidine dimers (CPDs) and pyrimidine-pyrimidone (6-4) photoproducts (6-4PP) trigger the Nucleotide Excision Repair (NER) pathway. NER involves more than 30 proteins, which choreograph a series of molecular events, including the unwinding of DNA, excision of nucleotides proximal to the lesion, re-synthesis of the excised bases, and ligation of the nascent DNA to the rest of the strand (Shah and He 2015). These events are accompanied by the upregulation of DDR checkpoint markers such as phosphor-H2AX and p53 Binding Protein 1 (53BP1) (Marti et al. 2006a; Huang et al. 2017).

While decades of research have revealed the biochemistry of NER and other DDR pathways in astonishing molecular details, most of these studies consider DDR an isolated process in which each single cell repairs its genome autonomously. In real life, radiations from sunlight or radiotherapy rarely cause direct DNA damages in more than a small subset of cells in the targeted tissue. In 1992, Nagasawa and Little reported that even when 1% of Chinese hamster ovary cells in a tissue culture dish were irradiated with plutonium-238 α-particles, DNA damages were detected in up to 30% of cells in the same dish (Nagasawa and Little 1992). This suggested that cells proximal to the irradiated cells could sustain DNA damages even without any exposure to radiation. The term Radiation-Induced Bystander Effect (RIBE) was coined to describe this phenomenon, in which genotoxicity such as micronuclei formation, DNA double-strand breaks, gene mutation, sister chromatid exchange, and oncogenic transformation are detected in unirradiated cells when their neighboring cells are irradiated (Freeman et al. 1993). These studies show that genotoxic factors are transmitted from irradiated cells to bystander cells via cell-cell interactions and soluble factors.

The crosstalk between irradiated and unirradiated cells appears bidirectional. Chen et al. reported that bystander unirradiated cells could release soluble factors that partially protect other cells from DNA damage (Chen et al. 2011). This intriguing phenomenon is termed Radiation-Induced Rescue Effect (RIRE) (Chen et al. 2011). Signaling molecules such as cyclic adenosine monophosphate (cAMP), Interleukin 6 (IL-6), and tumor necrosis factor-α (TNF-α) have been identified as “rescue signals” (Lam et al. 2015b, c; Kong et al. 2018). In response to these signals, the nuclear factor-kappa-light-chain-enhancer of activated B cell (NF-κB) pathway in the target cells is stimulated (Kong et al. 2018). Pathikonda et al. showed that poly (ADP-ribose) polymerase1 (PARP1) was upregulated in irradiated cells treated with rescue signals produced by bystander cells. Inhibition of NF-κB abolished this RIRE-mediated PARP1 upregulation, and inhibition of PARP1 suppressed the stimulation of the NF-κB pathway by RIRE signals (Pathikonda et al. 2020), suggesting NF-κB and PARP1 are engaged in a positive feedback loop in RIRE. The discovery of RIBE and RIRE underscores a complex intercellular signaling network in irradiated tissues that can modulate the outcome of radiation treatments. As the magnitude of both RIBE and RIRE depends on the precise radiation dosage (Buonanno et al. 2011; Yu 2019) and the relative number and cell types of irradiated and unirradiated cells (Azzam et al. 2013; Lam et al. 2015c), the net outcome of DNA damage and repair in an irradiated tissue are challenging to predict. The interplay between RIBE and RIRE has profound clinical implications, as it can lead to uncontrolled changes in genetic makeup in tissues beyond the irradiated area and the generation of protective cells that can compromise the effectiveness of subsequent sessions of radiation treatment.

Apart from a small number of in vivo studies involving partial irradiation of tumors by microbeams (Koturbash et al. 2007, 2008; Chai and Hei 2008; Ilnytskyy et al. 2009; Choi et al. 2011, 2012), our understanding of RIBE and RIRE are mostly from in-vitro studies based on co-culture and medium transfer experiments using immortalized cell lines cultured on flat plastic surfaces (Ballarini et al. 2002; Little 2006; Baskar 2010; Widel et al. 2012; Pereira et al. 2014; Desai et al. 2014; He et al. 2014; Wang et al. 2015; Lam et al. 2015a; Kong et al. 2018; Kirolikar et al. 2018; Lobachevsky et al. 2021). These setups do not reflect the anatomy of solid tumors, which are vascularised cell masses composed of cancerous and stromal cells encased in the extracellular matrix scaffold (Baker and Chen 2012). The deprivation of this microenvironment in these studies might prevent the fully appraisal of the physiological impact of RIBE and RIRE. The tumour microenvironment contains a complex assortment of immune cells, such as macrophages, neutrophils, natural killer cells, myeloid-derived suppressor cells (MDSCs), and mast cells (Hanahan and Coussens 2012; Albini et al. 2018). In particular, macrophages are pivotal to cancer homeostasis. As universal “transducers” (Moreira Lopes et al. 2020), macrophages constantly sample signals from their immediate surroundings and respond by secreting soluble factors that can directly affect the growth and proliferation of cancer cells or mobilize other immune components to regulate cell functions. Although collectively named as a single cell type, macrophages are highly plastic, malleable into heterogeneous phenotypes by environmental stimuli. Traditionally, two primary macrophage types are described (Viola et al. 2019). M1 macrophages are activated by ligands of toll-like receptors (TLR), such as lipopolysaccharides (LPS) or interferon-γ (IFN-γ). They are active in antigen presentation, expression of pro-inflammatory cytokines such as IL-12, IL-23, and TNF-α, and production of reactive radical species. M2 macrophages are induced by the activation of IL-4 and IL-13 receptors (M2a cells), immunoglobulin complexes with TLR agonists (M2b cells), or by IL-10, TGF-β, or glucocorticoids (M2c cells). It is generally believed that M1 macrophages are associated with inflammatory, microbicidal, and tumoricidal activities, whereas M2 macrophages resolve inflammation and promote healing by inducing cell proliferation (Italiani and Boraschi 2014). Though the classical M1/M2 paradigm lays an essential foundation for understanding macrophage functions, more refined gene expression analyses have revealed a far greater phenotypic plasticity in macrophages. For example, our group has recently discovered that magnesium ions (Mg^2+^) can polarise macrophages into a previously unknown phenotype during bone regeneration (Qiao et al. 2021). Furthermore, the tumour microenvironment can programme monocytes into a specific phenotype known as Tumour Associated Macrophages (TAMs), which play essential roles in immunosuppression, angiogenesis, extracellular matrix remodelling, and metastasis (Chanmee et al. 2014). In many studies, an in vitro model of TAM was generated by coculturing unpolarised macrophages with cancer cell lines or exposing macrophages to conditioned medium collected from cancer cell lines (Tjiu et al. 2009; Wei et al. 2019; Benner et al. 2019).

Macrophages can modulate DDR (Geiger-Maor et al. 2015). For instance, the DDR activity in breast cancer cells 4T1 was enhanced when co-cultured with TAMs (Soto et al. 2016), and the depletion of TAMs from Taxol-treated breast cancer cells enhanced DNA damages and apoptosis (Olson et al. 2017). Similarly, irradiated breast cancer cells MCF7 exhibited higher cell survival and lesser DNA damage when co-cultured with M2 macrophages (Lindström et al. 2017). M2 cells can promote the repair of Double-Strand Breaks (DSBs) in multiple myeloma cells through the non-homologous end-joining (NHEJ) process (Dong et al. 2019) and induce overexpression of Midkine (MDK), a transcriptional factor involved in the p53-DDR axis (Meng et al. 2019). Interestingly, the balance between M1 and TAM/M2 macrophages in tumors can be affected by irradiation according to the host’s genetic background. For example, macrophages in xenografts implanted in CBA/Ca mice exhibited M1 phenotypes upon irradiation, while those in C57BL/6 mice were predominantly M2 phenotypes (Coates et al. 2008). As a result, xenografts are radiosensitive when implanted in CBA/Ca mice but radioresistant in C57BL/6 mice. These studies indicate that the outcome of radiation on the survival of cancer cells in a tumour is strongly influenced by the dynamics of macrophage abundance and phenotypes [50].

Taken together, the response of individual cells in an irradiated tissue is influenced by the dynamic interplay of RIBE, RIRE, and immune cells. A clear understanding of how the interactions of these factors affect DDR is crucial to the accurate modeling of the radiation effect on healthy and pathological tissues. The magnitude of RIRE appears to be higher when non-transformed stromal cells, such as normal fibroblasts, were used as bystanders (Widel et al. 2012; Desai et al. 2014), suggesting that RIRE on cancer cells may be dependent on the tissue context. It is unknown how the soluble factors from macrophages in a tumour microenvironment may modulate the RIRE as bystander cells on another population of irradiated cancer cells. In this study, we examined the effect of macrophages of different phenotypes on the RIRE of MCF7 cells. Our data indicate a previously unknown role of macrophages in RIRE and imply that the phenotypes of macrophages in the tumor microenvironment can dramatically influence the efficiency of radiotherapeutics.

## Materials and methods

### Cell culture

MCF7 (mammary gland adenocarcinoma) and THP1 (human monocytic leukemia) were obtained from American Type Culture Collection (ATCC). These cell lines have been authenticated by Short Tandem Repeat (STR) profiling ([CSL STYLE ERROR: reference with no printed form.]). MCF7 were cultured in Dulbecco’s Modified Eagle Medium (DMEM; Gibco 10313021), and THP1 cells were cultured in Roswell Park Memorial Institute 1640 Medium (RPMI-1640; Gibco 11875093). Both media were supplemented with 10% fetal bovine serum (FBS) (Gibco 10270106), 1% GlutaMax™ (Gibco 35050061), and 1% penicillin-streptomycin (Gibco 15140122). Cells were incubated at 37°C in a humidified environment containing 5% CO2 and were passaged twice a week.

### Differentiation and polarisation of macrophages

THP1 cells were differentiated and polarised into four different subtypes as previously described (Chanput et al. 2013; Tedesco et al. 2018; Wei et al. 2019; Polumuri et al. 2021; Jian et al. 2021) and are shown in Fig. 1. To obtain differentiated macrophages (called M0 cells in this study), THP1 cells were incubated with 100ng/ml phorbol 12-myristate 13-acetate (PMA, Sigma Aldrich) for 48 h (Jian et al. 2021). To generate M1 macrophages, M0 cells were treated with 1ug/ul Lipopolysaccharide (LPS, Sigma) and 20ng/ml Interferon-gamma (IFN-γ, PeproTech) (Chanput et al. 2013; Tedesco et al. 2018). To generate M2 macrophages, M0 cells were treated with 40 ng/ml Interleukin 4 (IL-4, PeproTech) (Tedesco et al. 2018; Polumuri et al. 2021). After incubation for 48 h, the media were replaced by fresh culture medium (RPMI-1640 with 10% FBS). Phenotypes of polarised macrophages were confirmed by quantitating the mRNA levels of macrophage markers, including CD80 (Chen et al. 2020), CD86 (Kim et al. 2016), CD163 (Leung and Wong 2021), and CD206 (Kochiyama et al. 2019), using real-time PCR (see Table 1 for primer information). TAMs were established by co-culturing using 4×10^5^ M0 cells with 1.6×10^6^ MCF7 cells for 24 h, as previously described (Tjiu et al. 2009; Wei et al. 2019). To allow the convenient separation of the macrophages and MCF7 cells after coculture, these two cell types were cultured on separate coverslips in the same 100×20 mm Petri dishes, thus sharing the medium. After the co-culture, the coverslips carrying the macrophages were rinsed with fresh medium and transferred by forceps to other culture dishes for the subsequent steps of the experiment (see below).

**Fig. 1.**
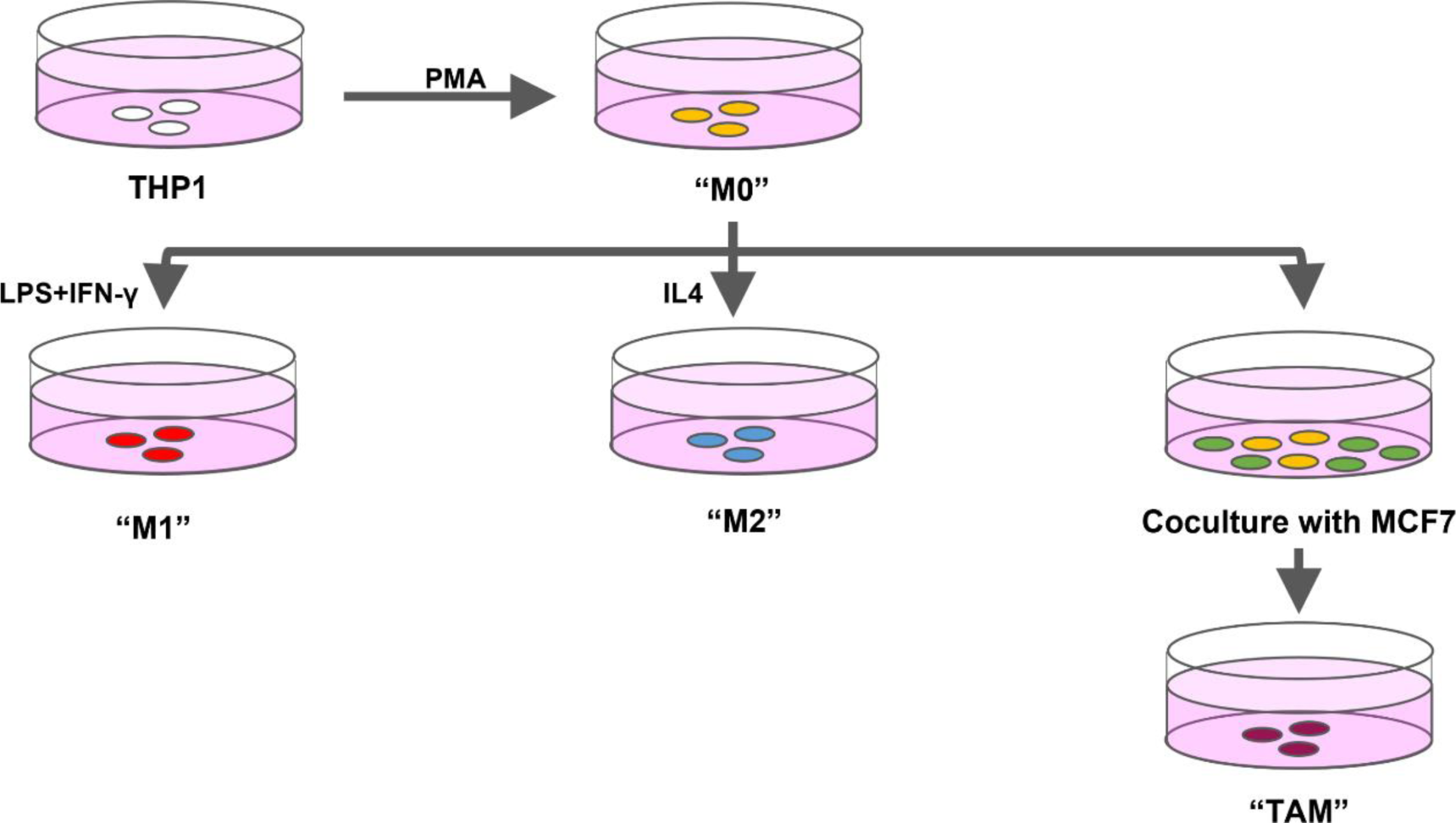
Schematic diagram showing the procedures for the differentiation and polarisation of THP1 cells into four different macrophage subtypes, namely, M0, M1, M2, and TAM

**Table 1.**
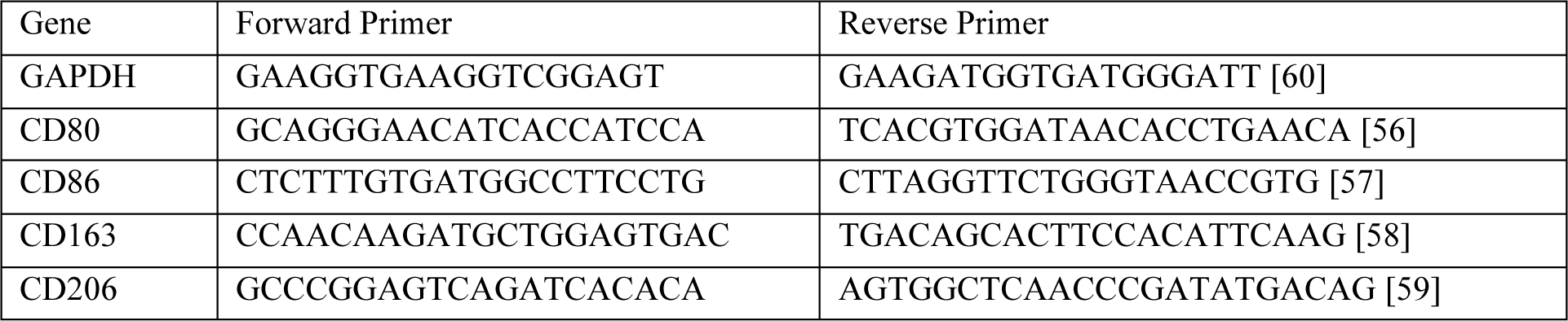
List of primer sequences for quantitative real-time polymerase chain reaction (q-RT PCR)

### Preparation of originator, bystander, and effector cells

In this study, we developed the procedure of bystander treatment (Fig. 2). To enable the convenient transfer and co-culture of various cell types, we seeded cancer cells or macrophages on 22×22 mm glass coverslips 24 hours before the experiment, at a density of 4×10^4^ cells per coverslip. These coverslips were cultured in 100×20 mm Petri dishes, which contained 10 ml of culture medium. Cells on each coverslip were thus exposed to the culture medium shared by other cells in the same dish. At different points of the experiments, these coverslips were transferred, using forceps, from one dish to another. For UV irradiation, glass coverslips containing MCF7 cells were washed with approximately three quick changes of PBS, with most of the PBS in the last wash drained off, thus minimizing the volume of liquid covering the cells during irradiation. A Stratalinker UV crosslinker 1800 (Stratagene) was then used to deliver UVC irradiation (254 nm) at a dosage of 50 J/m^2^. The duration of UV irradiation was approximately 2.5 seconds. In some experiments, the cells were mock-irradiated by being placed for the same duration in the UV crosslinker that remained switched off. The coverslips containing the irradiated or mock-irradiated cells (called “originator cells” in this study) were then transferred to a petri dish containing coverslips of another population of cells, called “bystander cells.” The bystander cells used in this study consisted of MCF7, M0, M1, M2, or TAM, depending on the experimental design. In each case, the ratio between the number of originator and bystander cells was 1:20. After co-culturing with the originator cells at 37°C for 2h, the coverslips of bystander cells were transferred to a new petri dish that contained 10mL of fresh culture medium (DMEM with 10% FBS), where they were cultured for a further 3h. Afterwards, the culture medium was collected from the dish (and labelled as “conditioned medium”). Meanwhile, another population of MCF7 cells (called “effector cells”) was UV-irradiated as previously described. After irradiation (or mock irradiation), the effector cells were cultured in 10mL of conditioned media obtained from bystander cells for a further 6 h before fixation and immunostaining.

**Fig. 2.**
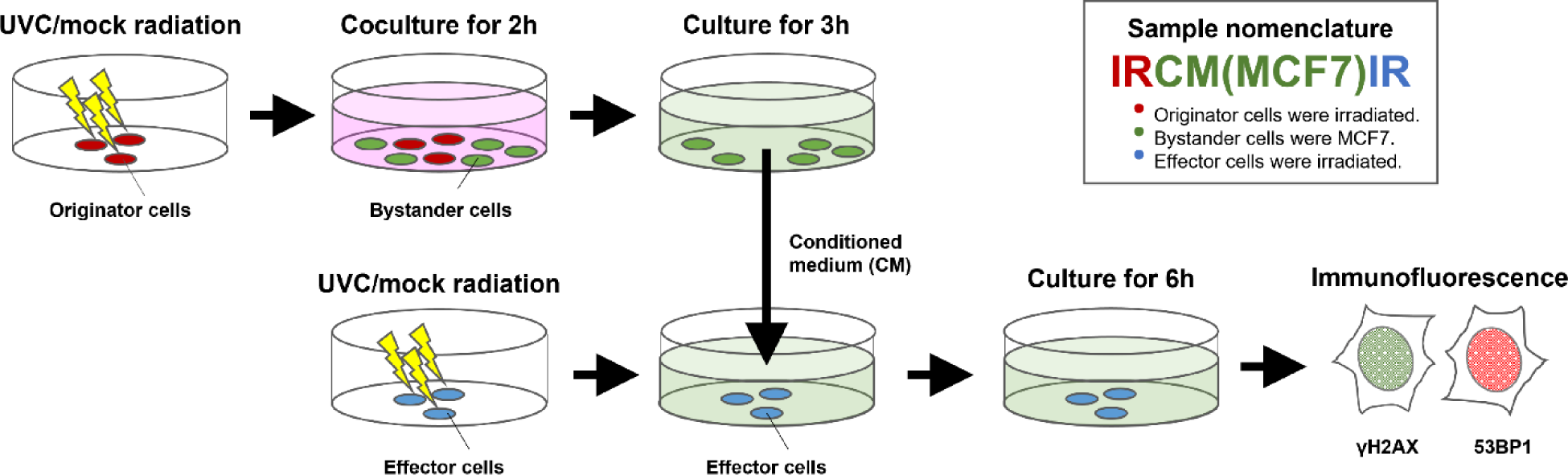
Schematic diagram showing the experimental design of this study. Inset illustrates the nomenclature of the samples. See text for explanations

### Immunofluorescence

Effector cells on coverslips were washed with phosphate-buffered saline (PBS) and fixed in 4% paraformaldehyde (PFA; Sigma) at room temperature (RT) for 10 min. The fixed cells were washed in PBS and then permeabilized using 0.25% Triton X-100 (Sigma) at RT for 10 min. Then, the cells were washed with PBS and blocked using 5% Bovine Serum Albumin (BSA, Sigma) for at least an hour before incubation with 1 ug/ml primary antibody anti-phospho-histone H2A.X (Ser139) (Millipore, 05-036) and primary anti-53BP1 (Abcam, ab175933) (1:250) at RT for 1h. The cells were washed in PBS and were incubated with 4ug/ml of secondary goat anti-mouse IgG (H+L) antibody conjugated with Alexa Fluor 633 (Invitrogen, A-21052) and 2ug/ml of secondary goat anti-rabbit IgG (H+L) antibody conjugated with Alexa Fluor 488 (Invitrogen, A-11008) at RT for 1h. Finally, the cells were washed with PBS, counterstained with Hoechst 33342 (1:10000) (Sigma), and then mounted on a glass slide using ProLong™ Glass Antifade Mountant (Invitrogen). The cells were imaged on a Laser Confocal Scanning Microscope (Leica SPE). The images were quantified using ImageJ software, and the corrected total nuclear fluorescence (CTNF) for each cell was measured using the formula: CTNF = Sum of the value of pixels in the selected nuclear area – (Nuclear area of selected cell × Mean fluorescence of background readings). A minimum of ∼40 cells from three experimental replicates for each sample were analysed.

### Real-time PCR analyses

RNA was isolated from 1×10^6^ THP1 cells after various treatments (see above) by using an RNAprep pure cell/bacteria kit (Tiangen Biotech 4992235) according to the manufacturer’s instructions and was quantified using NanoDrop One^C^ Spectrophotometer (Thermo Fisher Scientific). 1μg of isolated RNA was taken from each sample was then transcribed to cDNA using PrimeScript ™ RT reagent kit with gDNA Eraser (Takara Bio RR047B). According to the manufacturer’s instructions, real-time PCR was performed using TB Green® Premix Ex Taq™ (Tli RNaseH Plus) (Takara Bio RR420A). The signals were measured in the QuantStudio 3 Real-Time PCR System (Applied Biosystems). Glyceraldehyde 3-phosphate dehydrogenase (GAPDH) was used as a reference gene. Table 1 contains a list of primer sequences used for real-time PCR.

### Statistical analyses

Data here are represented as mean ± standard deviation (SD). The student’s t-test was employed to evaluate the differences between two sets of data (assuming unequal variances). The results corresponding to P < 0.05 were considered statistically significant for all the comparisons.

## Results

### Experimental design

In this study, we investigated how one population of UV-irradiated cells indirectly affected the level of DNA damage in another batch of UV-irradiated cells by acting on an intermediary, unirradiated population of cells and how this process was modulated by different macrophage subtypes. To achieve this, we spatially separated each cell population and sequentially exposed them to soluble factors released by another population (Fig. 2). Since this study involved many cell populations in a complicated medium transfer regimen, we coined the following terms to avoid confusion in reporting our results: “originators” are MCF7 cells irradiated with UV at the start of the experiment. After irradiation, these originators were co-cultured with a population of unirradiated cells called “bystanders” for 2h, during which the bystander cells were exposed to factors released by the originator cells. The bystanders used here were either MCF7, or a macrophage subtype (M0, M1, M2, or TAM), or a combination of both, e.g., M0+MCF7, M1+MCF7, etc. The bystanders were then transferred to a new dish and cultured for 3h. The medium collected during this 3h period is called the “Conditioned Medium” (CM). A fresh population of MCF7 cells, called “effectors,” was irradiated with UV. These effector cells were then exposed to the CM collected from the bystanders for 6h. We assessed the level of DNA damage in the effectors by nuclear immunofluorescence staining intensities of γH2AX and 53BP1. We believe this experimental design is an improvement of previous RIRE studies, in which DNA damages in irradiated cells cultured with bystanders or in bystander CM were compared to cells irradiated without bystanders (Chen et al. 2011; He et al. 2014; Lam et al. 2015c; Fu et al. 2016a). In these older studies, the irradiated cells in the test and control samples were often cultured in different cell densities or in media previously used by different number of cells, conditions that are known to affect DDR (Saeki et al. 1997; Long et al. 2003; Bar et al. 2004). In this study, the originators (irradiated cells used to stimulate bystanders) and the effectors (cells used to measure RIRE magnitude) were separated, and bystander cells cultured at the same density but treated with the CM of mock-irradiated originators were used as the control. As far as we know, this is the first study that adopts this experimental setup, in which any difference in DNA damages between the test and control samples would represent the effect specifically produced by bystander cells previously stimulated with irradiated cells.

To clearly identify the treatment regimen associated with each sample, we labeled the data of this study by a three-part code. The first part of the code signifies whether originator cells were UV- or mock-irradiated. UV-irradiation in the code is represented as “IR,” while the mock-irradiation is represented as “UIR.” The middle part signifies the identity of bystander cells. The third part signifies whether the effector cells were UV- or mock-irradiated. E.g. “IR-CM(MCF7)-IR” refers to the regimen in which the CM of MCF7 (bystanders) previously exposed to UV-irradiated MCF7 (originators) was used to treat another population of MCF7 (effectors) that has been UV-irradiated. “UIR-CM(MCF7)-IR” refers to the regimen in which the Originator cells (MCF7) were mock-irradiated. In order to establish the baseline levels of γH2AX and 53BP1 staining in MCF7 cells, we have also UV- or mock-irradiated previously unperturbed cells and then cultured them in fresh medium (FM) for 6h before immunostaining. These samples were labelled as “IR-FM” and “UIR-FM,” respectively.

### Radiation-induced rescue effect in UV-treated MCF7 cells

As expected, UV irradiation caused significant DNA damages in MCF7 cells, inducing a ∼6-fold and a ∼7-fold increase in the staining intensity of γH2AX and 53BP1, respectively (Fig. 3). In consistent with previous reports (Kruhlak et al. 2006; Marti et al. 2006b), the staining patterns of both γH2AX and 53BP1 were finely punctate throughout the nucleoplasm of UV-treated cells instead of large discrete nuclear foci detected in cells exposed to ionizing radiation. Therefore, we quantitated the levels of γH2AX and 53BP1 immunostaining by measuring the fluorescence intensity per unit nuclear area rather than counting the number of nuclear foci. As shown in Fig. 3, CM collected from bystander cells that have previously been co-cultured with UV-irradiated originator cells reduced γH2AX and 53BP1 staining in irradiated effector cells by 36% and 23%, respectively (IR-CM(MCF7)-IR vs. IR-FM, Fig. 3a, and 3b). CM from unperturbed MCF7 cells or MCF7 cells exposed to mock-irradiated cells could also cause a marginal reduction in γH2AX staining in the irradiated effectors (CM(MCF7)-IR and UIR-CM(MCF7)-IR, Fig. 3a, and 3b), though in a much smaller magnitude than IR-CM(MCF7)-IR. This result suggests that although the CM of MCF7 cells was inherently protective of UV-induced DNA damages, this protective effect was much more pronounced if the MCF7 cells were previously exposed to factors from DNA-damaged cells, consistent with RIRE previously described in cells stimulated by ionizing radiation (Pathikonda et al. 2020). As far as we know, this is the first unequivocal demonstration of RIRE as a rescue effect specifically generated by bystander cells previously exposed to factors from irradiated cells. Interestingly, CM from MCF7 cells, regardless of any previous treatment, could also reduce the baseline level of γH2AX and 53BP1 staining in unirradiated cells (Supplementary Fig. 1b and 1c, see online resource).

**Fig. 3.**
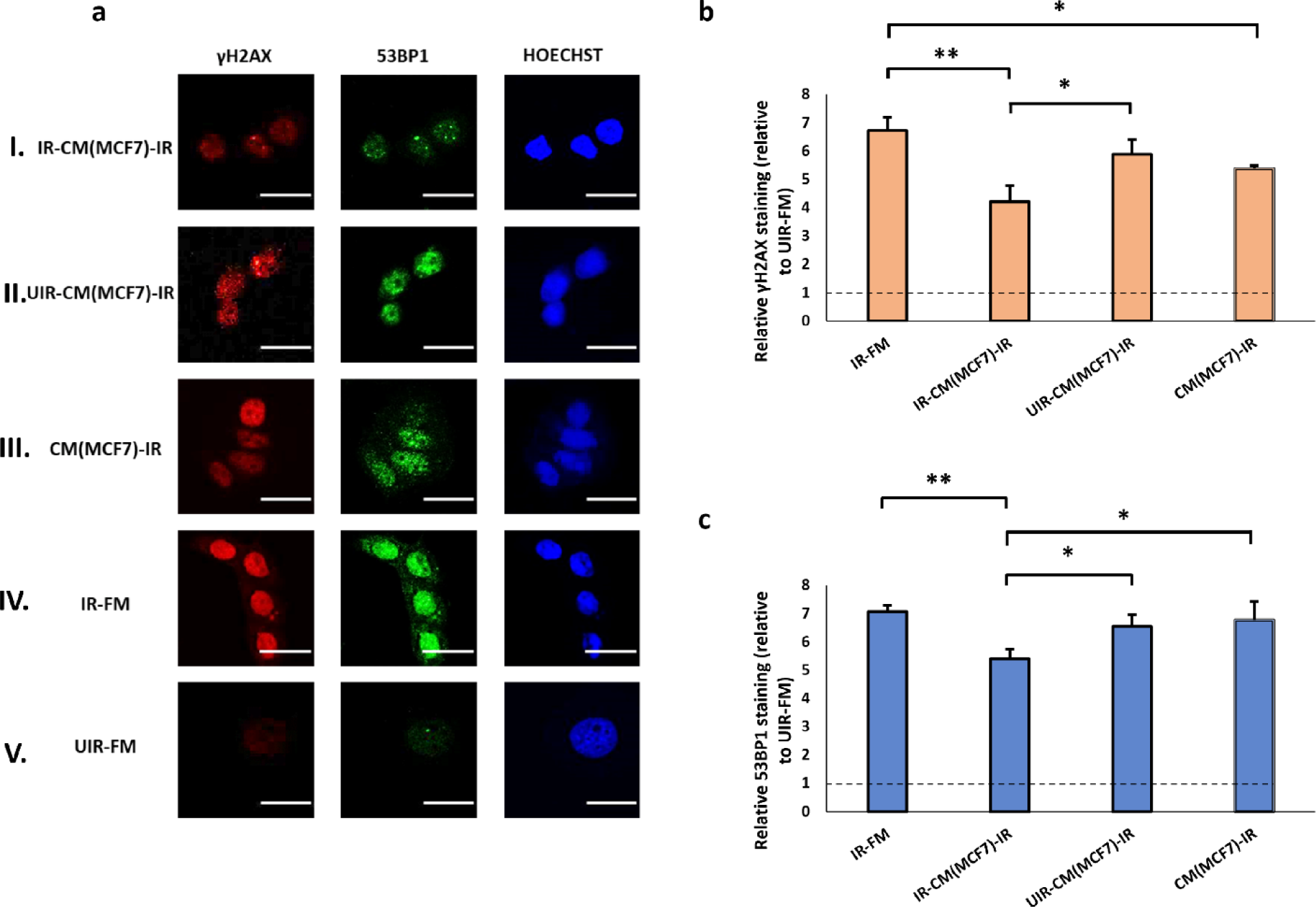
(a) Representative images from immunofluorescence staining with anti-phospho-histone H2A.X and anti-53BP1 antibodies in MCF7 cells post-UVC irradiation. CM experiments with MCF7 cells as Bystanders were carried out in the following conditions, (I) IR-CM(MCF7)-IR, (II) UIR-CM(MCF7)-IR, (III) CM(MCF7)-IR. Control experiments included (IV) IR-FM and (V) UIR-FM. Scale bar = 25 μm. (b) Graph representing relative γH2AX staining relative to that in UIR-FM cells, which is set as the baseline (dotted lines) for these conditions. (c) Graph representing relative 53BP1 staining (relative to UIR-FM) for these conditions. * *P* < 0.05; ** *P* < 0.01 and error bars represent mean ± SD

### Soluble factors from unpolarised macrophages reduce DNA damages in UV-radiated cells

We then asked whether RIRE occurred when macrophages were used as bystanders. In this study, we used macrophages differentiated from monocytic cell line THP-1 as our model (Auwerx 1991). We observed a pronounced rescue effect from the CM of differentiated THP-1 cells previously exposed to irradiated MCF7 cells, causing a 2.1 and 1.8-fold decrease in γH2AX and 53BP1 staining in irradiated effector cells (IR-CM(M0)-IR, Fig. 4a, and 4b). This effect was greater than that of bystander MCF7 cells (Fig. 3). However, rescue effects of similar magnitude were also detected for the macrophage CM without pre-exposure to UV-damaged cells (UIR-CM(M0)-IR and CM(M0)-IR, Fig. 4). We also tested the effect of macrophage CM on the baseline levels of γH2AX and 53BP1 staining in unirradiated MCF7 cells. Unlike MCF7 cells, macrophage CM did not cause significant changes in γH2AX and 53BP1 staining unirradiated cells (Supplementary Fig. 2b and 2c, see online resource). These results suggested that the rescue effect of macrophages on DNA damages appeared to be distinct from that of cancer cells. Factors from macrophages could inherently mitigate DNA damages in UV-irradiated MCF7 cells regardless of whether these macrophages have previously been primed by UV-damaged cells, whereas the protective effects of factors from MCF7 cells were more pronounced when exposed to DNA-damaged originators.

**Fig. 4.**
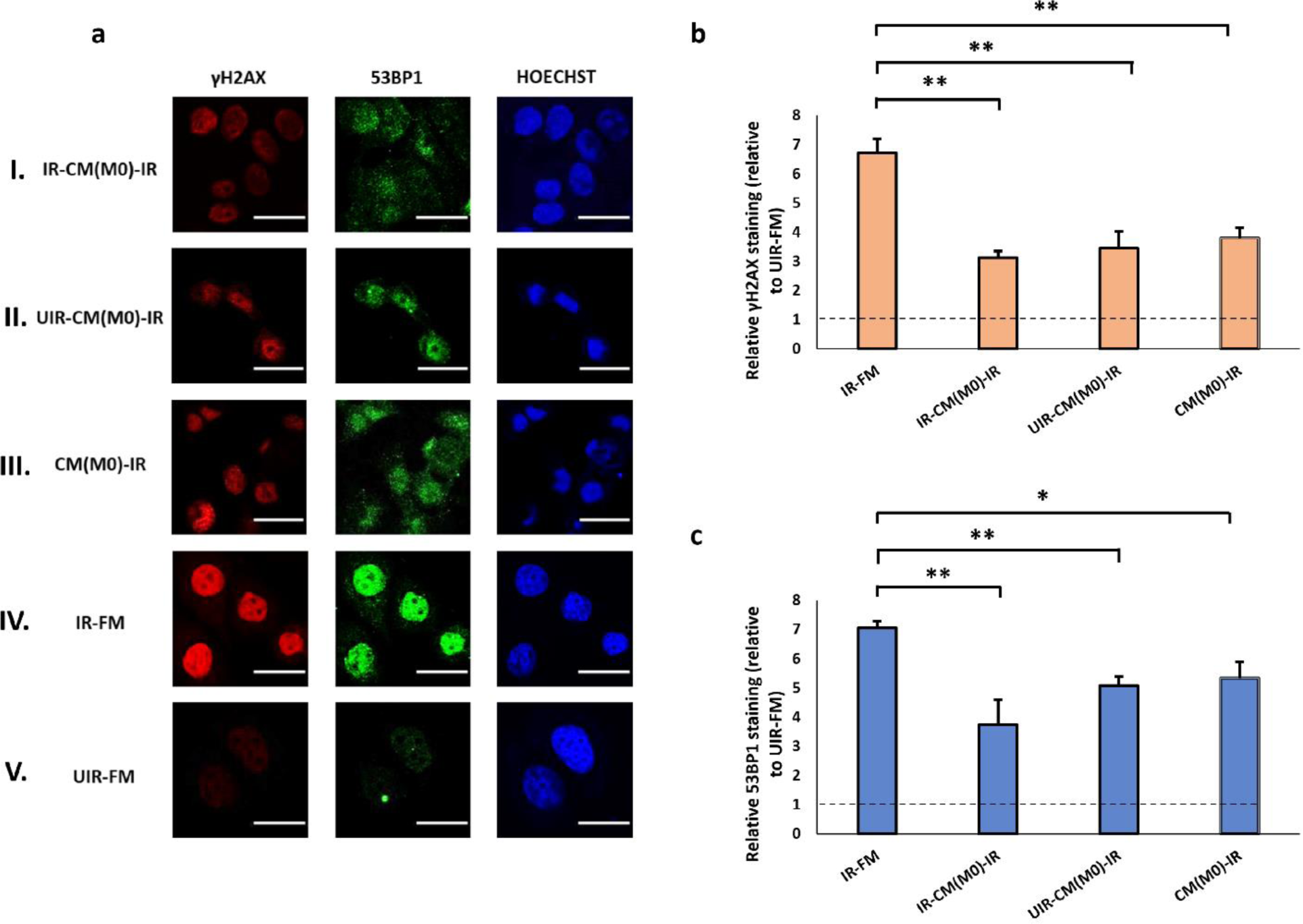
(a) Representative images from immunofluorescence staining with anti-phospho-histone H2A.X and anti-53BP1 antibodies in MCF7 cells post-UVC irradiation. CM experiments with M0 cells as Bystanders were carried out in the following conditions, (I) IR-CM(M0)-IR, (II) UIR-CM(M0)-IR, (III) CM(M0)-IR. Control experiments included (IV) IR-FM and (V) UIR-FM. Scale bar = 25 μm. (b) Graph representing relative γH2AX staining (linear values relative to UIR-FM) for these conditions. (c) Graph representing relative 53BP1 staining (linear values relative to UIR-FM) for these conditions. * *P* < 0.05; ** *P* < 0.01 and error bars represent mean ± SD

### Radiation-induced rescue effects of macrophages depend on polarisation phenotypes

Next, we asked if the polarisation of macrophages could modulate its protective effect on DNA damages. We induced M1 and M2 polarisation by treating differentiated THP-1 cells with LPS/IFN-γ and IL-4, respectively. RT-PCR analyses confirmed the upregulation of M1 markers CD80 and CD86 (Nielsen et al. 2020) and M2 markers CD163 and CD206 (Hedbrant et al. 2015) after respective treatments (Supplementary Fig. 3, see online resource). Apart from M1 and M2 cells, we also generated TAM by co-culturing differentiated THP-1 cells with MCF7 cells (Wei et al. 2019).

We tested the effect of the CM from these polarised macrophages, collected after exposure to UV-or mock-irradiated originator cells, on γH2AX and 53BP1 levels in MCF7 cells. M1 CM caused minor or insignificant changes in the γH2AX and 53BP1 staining in irradiated cells regardless of whether the M1 cells were primed by UV-damaged cells or not (Fig. 5b and 5c). Similarly, M1 CM caused insignificant changes in the 53BP1 staining in irradiated cells, regardless of whether the M1 cells have been primed by UV-damaged cells or not (Fig. 5c). The effect of M1 CM on the baseline levels of γH2AX and 53BP1 in unirradiated MCF7 cells was also marginal (Supplementary Fig. 4b and 4c, see online resource). By contrast, the CM of M2 and TAM strongly reduced γH2AX and 53BP1 levels in irradiated MCF7 cells (Fig. 6 and 7), in consistent with the established pro-survival roles of these two macrophage subtypes (Genin et al. 2015; Chen et al. 2019). The suppression of DNA damages was the most pronounced for the CM of M2 and TAM previously exposed UV-irradiated originators (IR-CM(M2)-IR, Fig. 6; and IR-CM(TAM)-IR, Fig. 7). This suggests the protective effect of M2 and TAM could be further enhanced by RIRE, unlike that of M0 cells. In particular, the RIRE of M2 cells nearly reduced the γH2AX and 53BP1 staining of the irradiated effector cells to the baseline level (UIR-FM), suggesting that CM from bystander M2 or TAM could almost fully protect MCF7 cells from UV damage. Interestingly, pre-exposure to UV-damaged originator cells did not significantly increase the effect of M2 and TAM CM on the baseline γH2AX and 53BP levels in unirradiated MCF7 cells (Supplementary Fig. 5b, 5c, 6b, and 6c, see online resource).

**Fig. 5.**
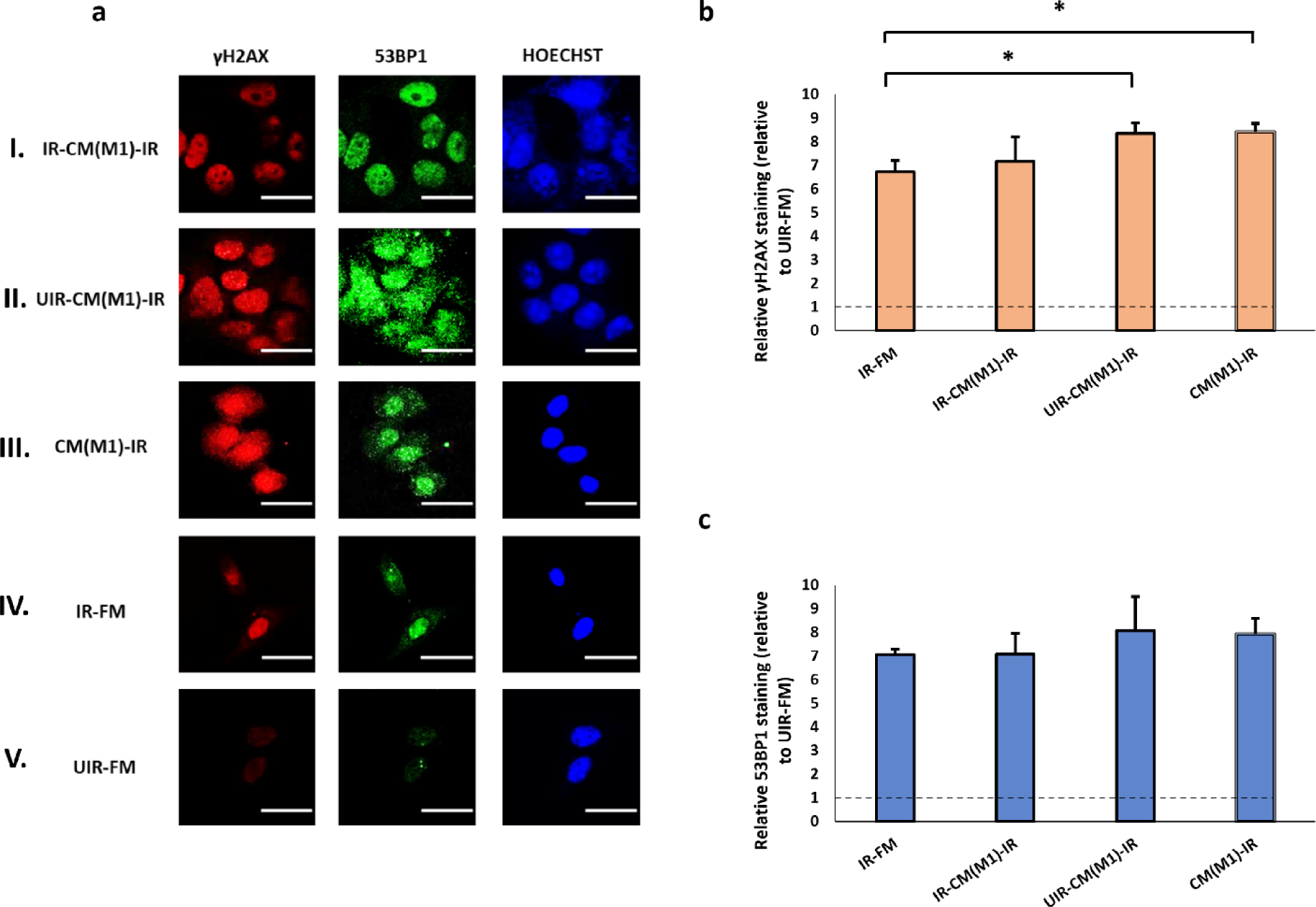
(a) Representative images from immunofluorescence staining with anti-phospho-histone H2A.X and anti-53BP1 antibodies in MCF7 cells post-UVC irradiation. CM experiments with M1 cells as Bystanders were carried out in the following conditions, (I) IR-CM(M1)-IR, (II) UIR-CM(M1)-IR, (III) CM(M1)-IR. Control experiments included (IV) IR-FM and (V) UIR-FM. Scale bar = 25 μm. (b) Graph representing relative γH2AX staining (linear values relative to UIR-FM) for these conditions. (c) Graph representing relative 53BP1 staining (linear values relative to UIR-FM) for these conditions. * *P* < 0.05; ** *P* < 0.01 and error bars represent mean ± SD

**Fig. 6.**
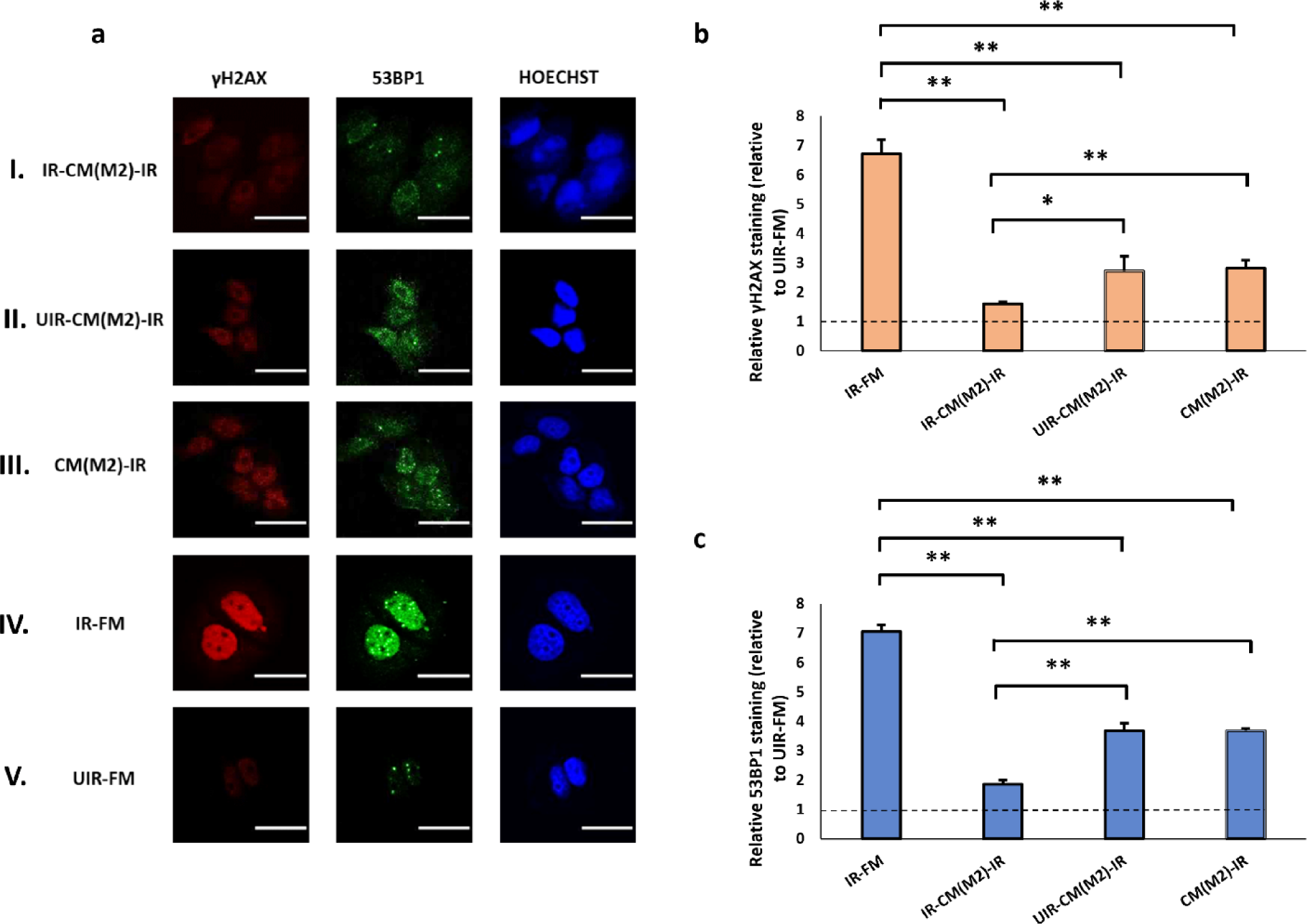
(a) Representative images from immunofluorescence staining with anti-phospho-histone H2A.X and anti-53BP1 antibodies in MCF7 cells post-UVC irradiation. CM experiments with M2 cells as Bystanders were carried out in the following conditions, (I) IR-CM(M2)-IR, (II) UIR-CM(M2)-IR, (III) CM(M2)-IR. Control experiments included (IV) IR-FM and (V) UIR-FM. Scale bar = 25 μm. (b) Graph representing relative γH2AX staining (linear values relative to UIR-FM) for these conditions. (c) Graph representing relative 53BP1 staining (linear values relative to UIR-FM) for these conditions. * *P* < 0.05; ** *P* < 0.01 and error bars represent mean ± SD

**Fig. 7.**
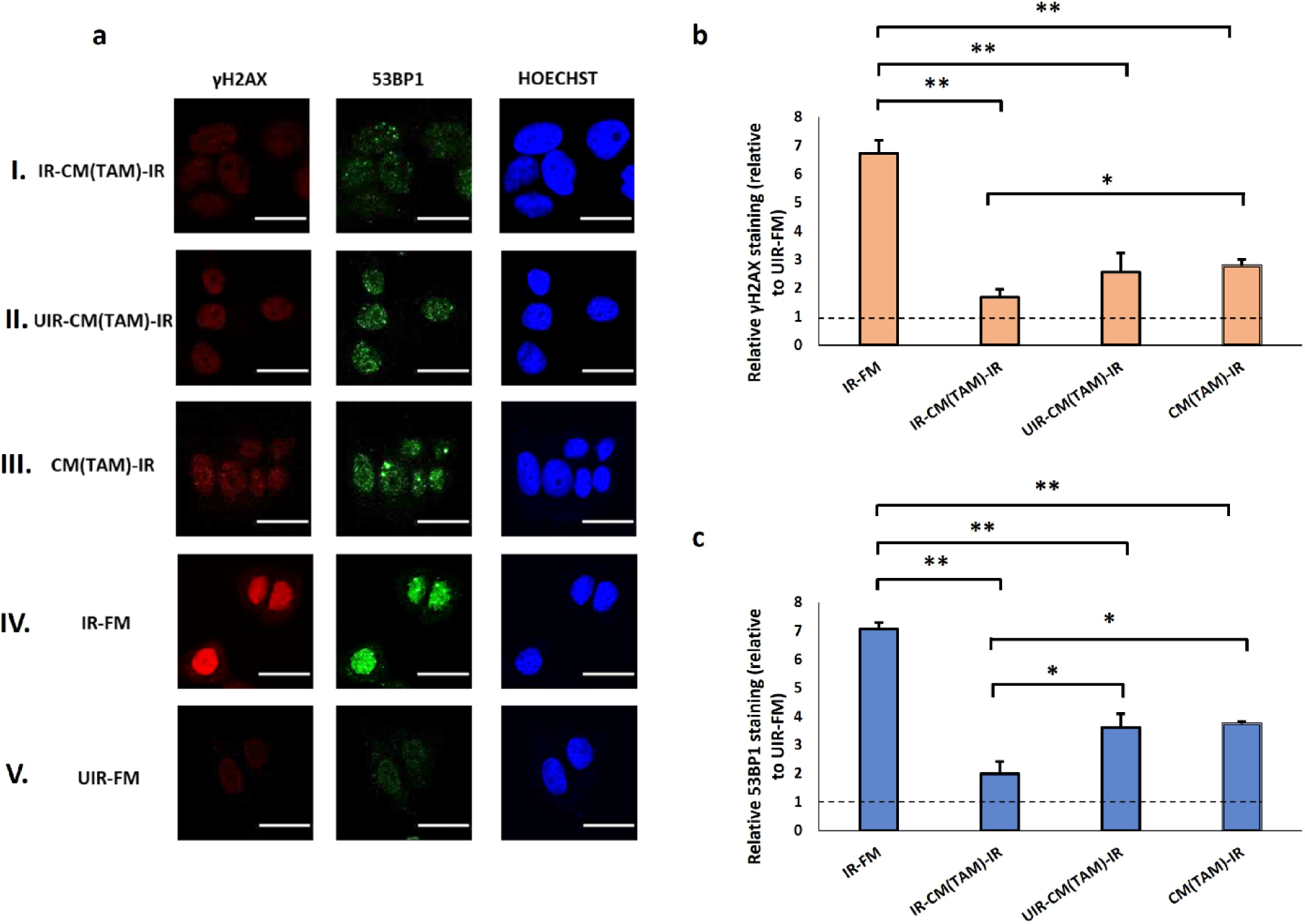
(a) Representative images from immunofluorescence staining with anti-phospho-histone H2A.X and anti-53BP1 antibodies in MCF7 cells post-UVC irradiation. CM experiments with TAMs as Bystanders were carried out in the following conditions, (I) IR-CM(TAM)-IR, (II) UIR-CM(TAM)-IR, (III) CM(TAM)-IR. Control experiments included (IV) IR-FM and (V) UIR-FM. Scale bar = 25 μm. (b) Graph representing relative γH2AX staining (linear values relative to UIR-FM) for these conditions. (c) Graph representing relative 53BP1 staining (linear values relative to UIR-FM) for these conditions. * *P* < 0.05; ** *P* < 0.01 and error bars represent mean ± SD

### Radiation-induced rescue effect of MCF7 cells is modulated by macrophages

The experiments reported above investigated RIRE in which the bystander cells were either MCF7 or macrophages. In a tumour microenvironment, immune and tumour cells intermingled in close proximity. We, therefore, considered a scenario in which a co-culture of MCF7 cells and macrophages (in a 1:1 ratio) acted in combination as bystander cells. As shown in Fig. 8, the co-culture of MCF7 and M1 cells in the bystander population moderately reduced γH2AX staining in UV-irradiated MCF7 cells, as compared to the use of M1 as bystander cells alone (IR-CM(M1)-IR vs. IR-CM(M1+MCF7)-IR, Fig. 8). However, the presence of MCF7 cells in the bystander co-culture of M0, M2, and TAM did not critically affect the magnitude of RIRE (Fig. 8). In fact, MCF7 alone as bystanders did not reduce the γH2AX and 53BP1 staining as much as the above-mentioned bystanders M0, M2, and TAM with or without MCF7. These data suggest that the effect of macrophages dominated the overall RIRE induced by a combination of MCF7 cells and macrophages. These results highlight the importance of macrophages in the study of RIRE of cancer cells.

**Fig. 8.**
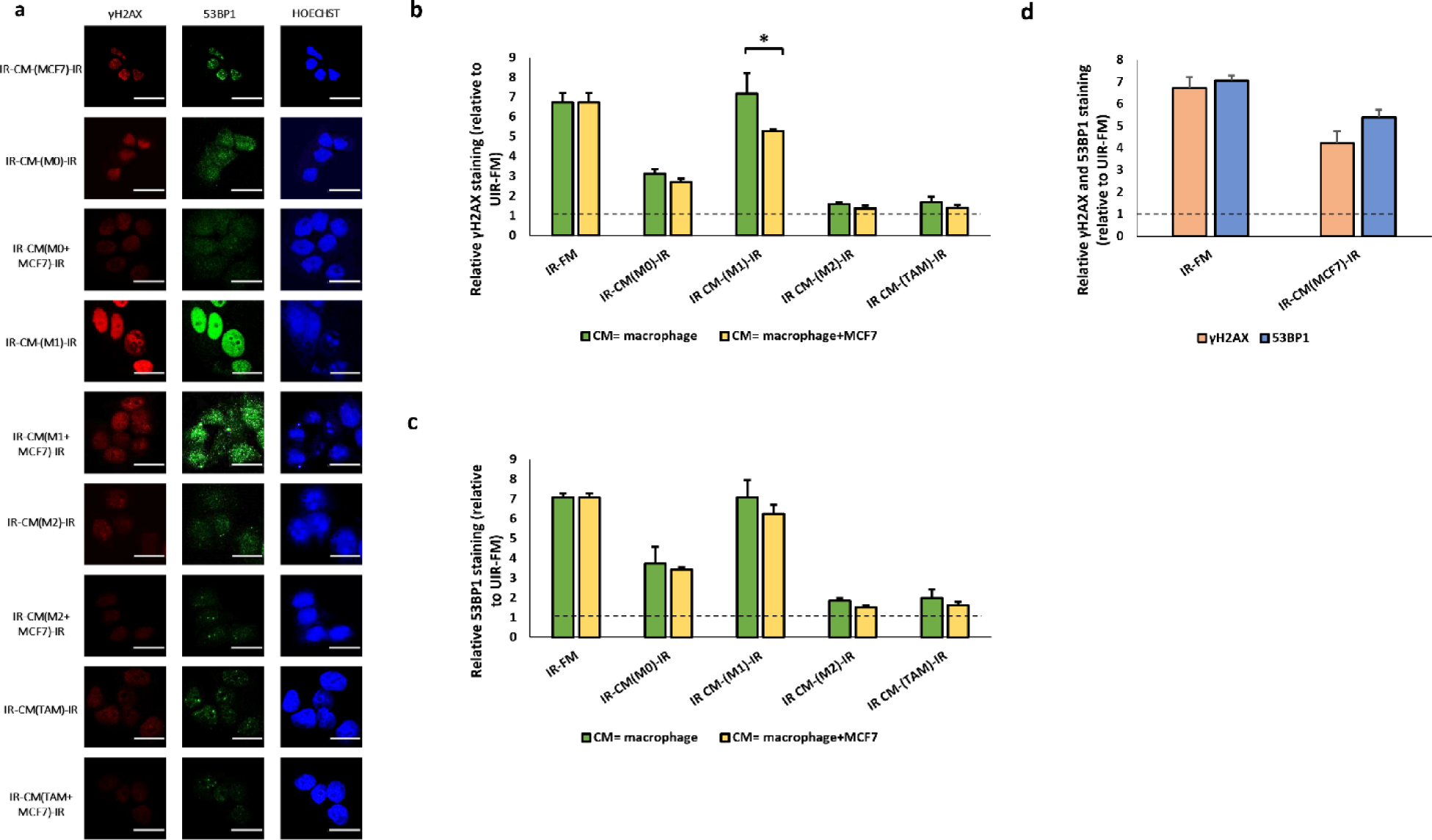
The Fig. represents the data obtained from MCF7 cells treated with CM obtained from Bystanders of MCF7 and different macrophage subtypes (M0, M1, M2, and TAMs) with or without MCF7, which were previously co-cultured with Originators. (a) Representative images from immunofluorescence staining with anti-phospho-histone H2A.X and anti-53BP1 antibodies in MCF7 cells post-UVC irradiation. Scale bar = 25 μm. * *P* < 0.05; ** *P* < 0.01 and error bars represent mean ± SD. (b) The graph represents relative γH2AX staining (linear values relative to UIR-FM) for IR-CM-IR (macrophage) and IR-CM-IR (macrophage and MCF7). (c) The graph represents the average fluorescent intensities of γH2AX for the above-said conditions. (d) The graph represents relative γH2AX and 53BP1 staining (linear values relative to UIR-FM) for IR-CM-IR (MCF7)

## Discussion

This study examines the effect of macrophages on RIRE on UV-induced DNA damages in MCF7 cells. Our carefully controlled experiments allowed the separation of the effects of “naïve” and “stimulated” bystanders on DNA damages of irradiated cancer cells. We defined naïve bystanders as cells that have not been exposed to irradiated cells, whereas stimulated bystanders are previously exposed to factors from irradiated cells. All previous studies on RIRE compared the levels of DNA damage in cells irradiated with and without bystanders. Those studies could not determine to what extent the observed reduction of DNA damages was actually caused by stimulated bystanders and whether naïve bystander cells could confer some protection on irradiated cells. Moreover, the modulatory effect of cell density on DDR was not controlled. We addressed these limitations in the present study. Here, we demonstrated that, with the exception of unpolarised macrophages (M0 cells), CM from stimulated bystanders offered a higher level of protection against UV-induced DNA damages in MCF7 cells than naïve bystanders. Thus, RIRE is a specific and active response of stimulated bystander cells, possibly as a physiologically significant outcome of RIBE. We propose to name the protective effect provided specifically by stimulated bystanders the “active RIRE” and the effect that is common to both naïve and stimulated bystanders the “basal RIRE.”

Our data showed that active RIRE was not evident when unpolarised THP1-derived macrophages (M0) were used as bystander cells. In consistent with the established pro-survival roles of M2/TAM cells, we observed that soluble factors from M2 and TAM were highly protective of UV-irradiated MCF7 cells from DNA damages. The protective effect of M2 and TAM could be further enhanced by prior exposure to factors from UV-damaged cells, suggesting that macrophages became capable of conferring active RIRE upon M2/TAM polarisation. We also demonstrated that the presence of M1 macrophages as bystanders could reduce the RIRE on MCF7 cells. Hence, our data showed that the relative contribution of the active and basal RIRE in macrophages could be altered by polarisation. Non-cancerous cell lines such as hepatocytes and fibroblasts have been reported to generate RIRE (Chen et al. 2011; Widel et al. 2012; Pereira et al. 2014; He et al. 2014; Lam et al. 2015a; Fu et al. 2016b), but it was not previously known that RIRE of a particular cell type can be adjusted by changes in environmental conditions.

Although discovered for more than a past decade, all studies on RIRE have so far been associated with genotoxicity caused by ionizing radiations (Yu 2019). Diverse genotoxic stresses, including non-ionizing radiation, chemotherapeutics, heavy metals, and nanoparticles, have been demonstrated to induce RIBE (Verma and Tiku 2017). It is not known whether these stresses can also invoke RIRE. In this study, we demonstrated the occurrence of RIRE in UV-damaged MCF7 cells. Future investigations will be focused on whether the UV-RIRE involves the same rescue factors and molecular mechanisms, such as the NF-κB and PARP1 pathways, as in ionizing-RIRE (Pathikonda et al. 2020).

Our data showed that active RIRE was generated by M2 and TAM but not M0 and M1, suggesting that M2 and TAM are responsive to factors from irradiated originators. Irradiated cells affects neighbouring bystander cells by secreting “bystander signalling factors” such as IL-33, 1L-6, IL-8, TNF-α, TGF-β, and COX-2 (K. Hei et al. 2011; Desai et al. 2013). These factors are also known to activate M2 and TAM but are not required for M1 polarisation (Na et al. 2013; Casella et al. 2016; Zhang et al. 2016; Xiao et al. 2018; Fournié and Poupot 2018). Hence, it is possible that the phenotypic characteristics of M2 and TAM were enhanced when these cells were exposed, as bystanders, to the CM of irradiated originators. Furthermore, the NF-κB and PARP1 pathways are known to be activated in the originators (Lam et al. 2015c; Pathikonda et al. 2020) and are also involved in the commitment of macrophages to their M2 and TAM phenotypes (Hagemann et al. 2008; Sobczak et al. 2020). It is possible that, due to the pre-activation of NF-κB and PARP1, M2 and TAM are more responsive to stimulations from the irradiated originators.

In a tumour environment, macrophages and cancer cells intermingle in a close proximity. We investigated the scenario where macrophages and MCF7 cells were mixed in a bystander culture and how this mixture modulated RIRE. Our data showed that M1 cells could reduce the RIRE generated by MCF7 cells, in consistent with the role of M1 macrophages in aggravating DNA damages (Genard et al. 2018). These results are consistent with a previous observation that bystander U937 macrophages, which might have exhibit M1 characteristics (Widel et al. 2012), reduce the survival of α particle-irradiated human bronchial epithelial (Beas-2B) cells (Fu et al. 2016b). Our observation that M1 could potentially enhance the effectiveness of radiation therapy by removing RIRE adds one more therapeutic advantage for the raising of M1 abundance in tumours (Travers et al. 2019).

We observed that although M2, TAM, and MCF7 could individually confer active RIRE, the mixture of M2 or TAM with MCF7 as bystander cells did not generate a stronger RIRE. For example, CM of MCF7 and M2 cells, primed with UV-irradiated originators, were found to reduce γH2AX staining in irradiated effectors by 37% and 76%, respectively. But when mixed in a 1:1 ratio in the bystander culture, the overall reduction of γH2AX staining was only 79%. Similarly, the detected RIRE of TAM and MCF7 mixture was 79%, whereas TAM alone could induce a reduction of 75% already. It appears that the RIRE induced by a mixture of macrophages and cancer cells was not the direct summation by the individual effect from each of these two cell types. It is possible that the stronger RIRE of M2 or TAM subsumed the weaker effect of MCF7, or the two cell populations interact in a complex, nonlinear manner. A better understanding of the molecular mechanism underlying RIRE will be necessary to explain our observations. The dominating contribution of macrophages to RIRE even in the presence of 50% MCF7 cells in the bystander population underscores the importance of including macrophages in all RIRE studies. As the magnitude of RIRE is so dependent on the type of macrophages present in the bystander cell population, the physiological relevance of previous RIRE studies based on homogenous cancer cell lines without considering the contribution of macrophages may need to be revised. Future investigations will also involve the varying of ratios and spatial juxtaposition, in 3D cultures (Kunz-Schughart et al. 2004) of macrophage and cancer cells. Also, this study mainly focused on γH2AX and 53BP1, two well-known markers of genotoxicity. It will be interesting to investigate the expression level of other components of the NER pathway under these conditions.

In conclusion, the results of this study implicate that the macrophages in a tumour microenvironment affect the effect of non-ionizing radiation or other genotoxic therapies on cancer cells by modulating RIRE. The results also show that RIRE is not specific to the genotoxicity caused by DSBs but can occur for UVC-induced DNA lesions or photoproducts. Future investigations on different types of cancers will reveal if the observed effects were specific to breast cancer alone or applicable to other cancer types. It will also be interesting to elucidate the roles of other stromal components of the tumour microenvironment, such as other immune cells and tumour-associated fibroblasts, in RIRE.

## Compliance with ethical standards Conflict of interests

The authors declare no conflict of interests.

## Ethical approval

Not applicable.

## Informed consent

Not applicable.

