## Supplementary figures for "Radiation-induced rescue effect on human breast carcinoma cells is regulated by macrophages"

[3]School of Applied Sciences. University of Huddersfield, Huddersfield, UK.

### Supplementary information

The following are supplementary data to this article:

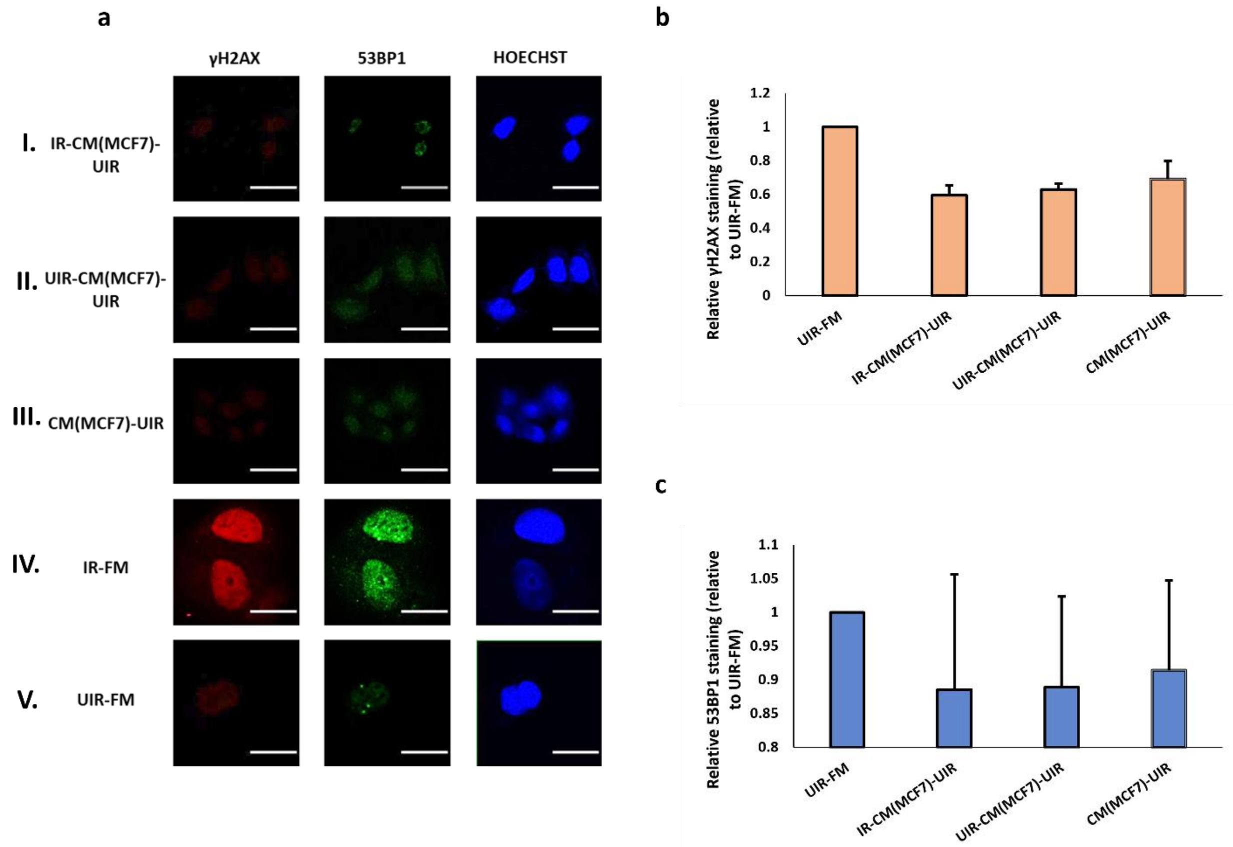


**Supplementary Fig. 1** (a) Representative images from immunofluorescence staining with anti-phospho-histone H2A.X and anti-53BP1 antibodies in MCF7 cells post-UVC mock-irradiation. CM experiments with MCF7 cells as Bystanders were carried out in the following conditions, (I) IR-CM(MCF7)-UIR, (II) UIR-CM(MCF7)-UIR, (III) CM(MCF7)-UIR. Control experiments included (IV) IR-FM and (V) UIR-FM. Scale bar = 25 µm. (b) Graph representing relative γH2AX staining (linear values relative to UIR-FM) for these conditions. (c) Graph representing relative 53BP1 staining (linear values relative to UIR-FM) for these conditions. * $P < 0.05$; ** $P < 0.01$ and error bars represent mean ± SD

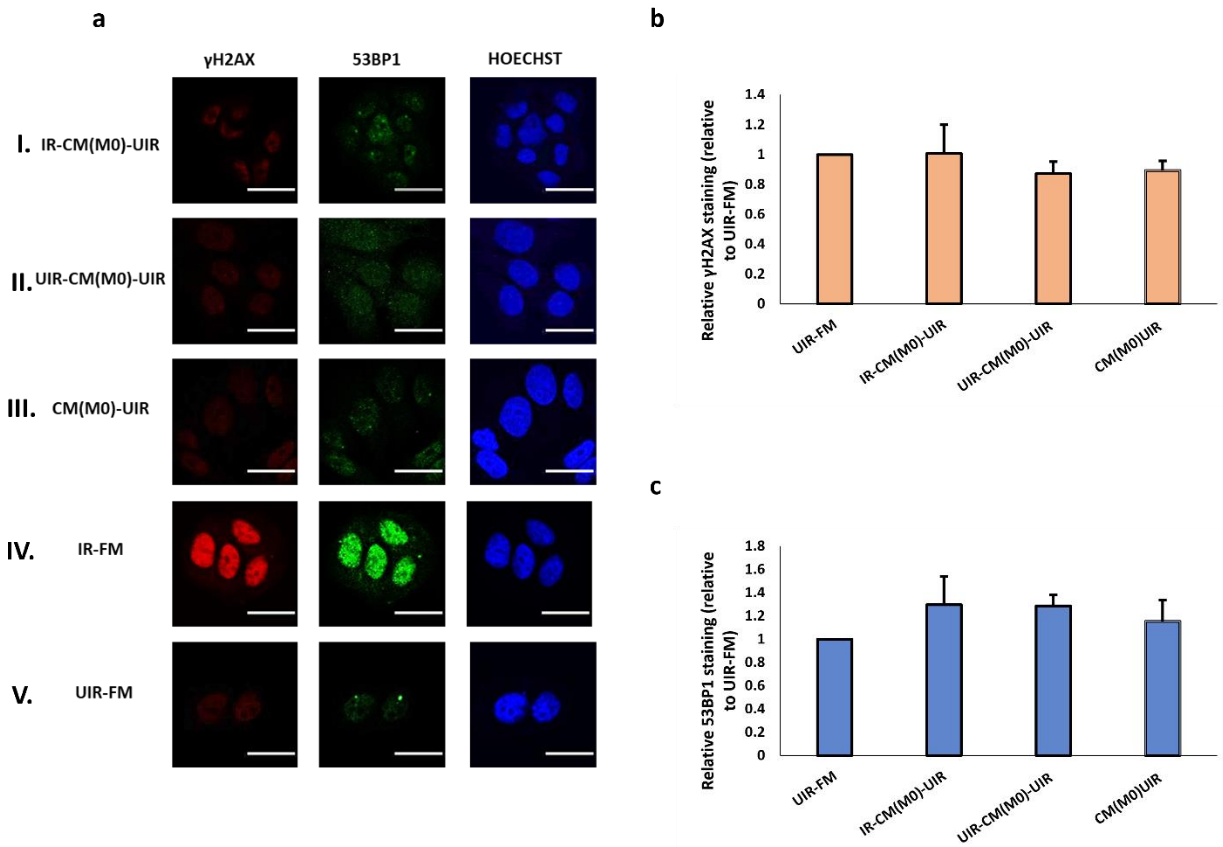


**Supplementary Fig. 2** (a) Representative images from immunofluorescence staining with anti-phospho-histone H2A.X and anti-53BP1 antibodies in MCF7 cells post-UVC mock-irradiation. CM experiments with M0 cells as Bystanders were carried out in the following conditions, (I) IR-CM(M0)-UIR, (II) UIR-CM(M0)-UIR, (III) CM(M0)-UIR. Control experiments included (IV) IR-FM and (V) UIR-FM. Scale bar = 25 μm. (b) Graph representing relative γH2AX staining (linear values relative to UIR-FM) for these conditions. (c) Graph representing relative 53BP1 staining (linear values relative to UIR-FM) for these conditions. * $P < 0.05$; ** $P < 0.01$ and error bars represent mean ± SD

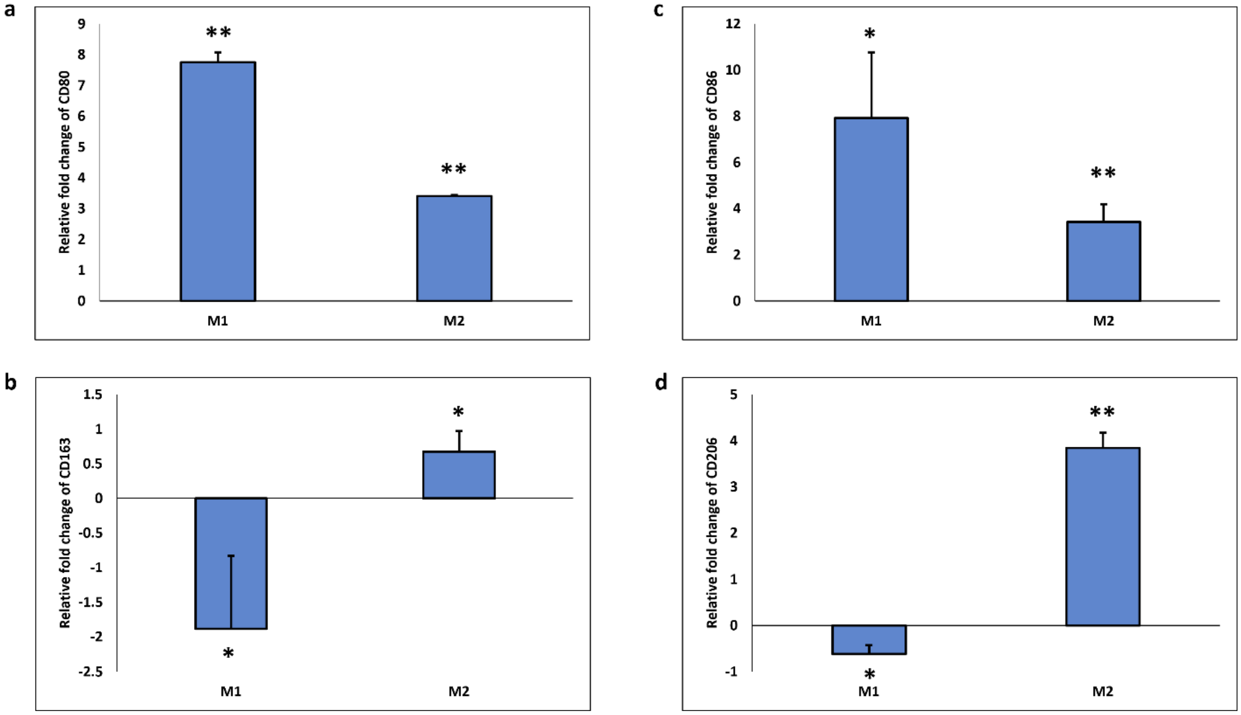


**Supplementary Fig. 3** Relative mRNA expression levels of M1 and M2 macrophage activation markers, (A) CD80, (B) CD163, (C) CD86, and (D) CD206 determined to check the polarities in M1 and M2 phenotypes against the control (M0). *$P < 0.05$; **$P < 0.01$ and error bars represent SD

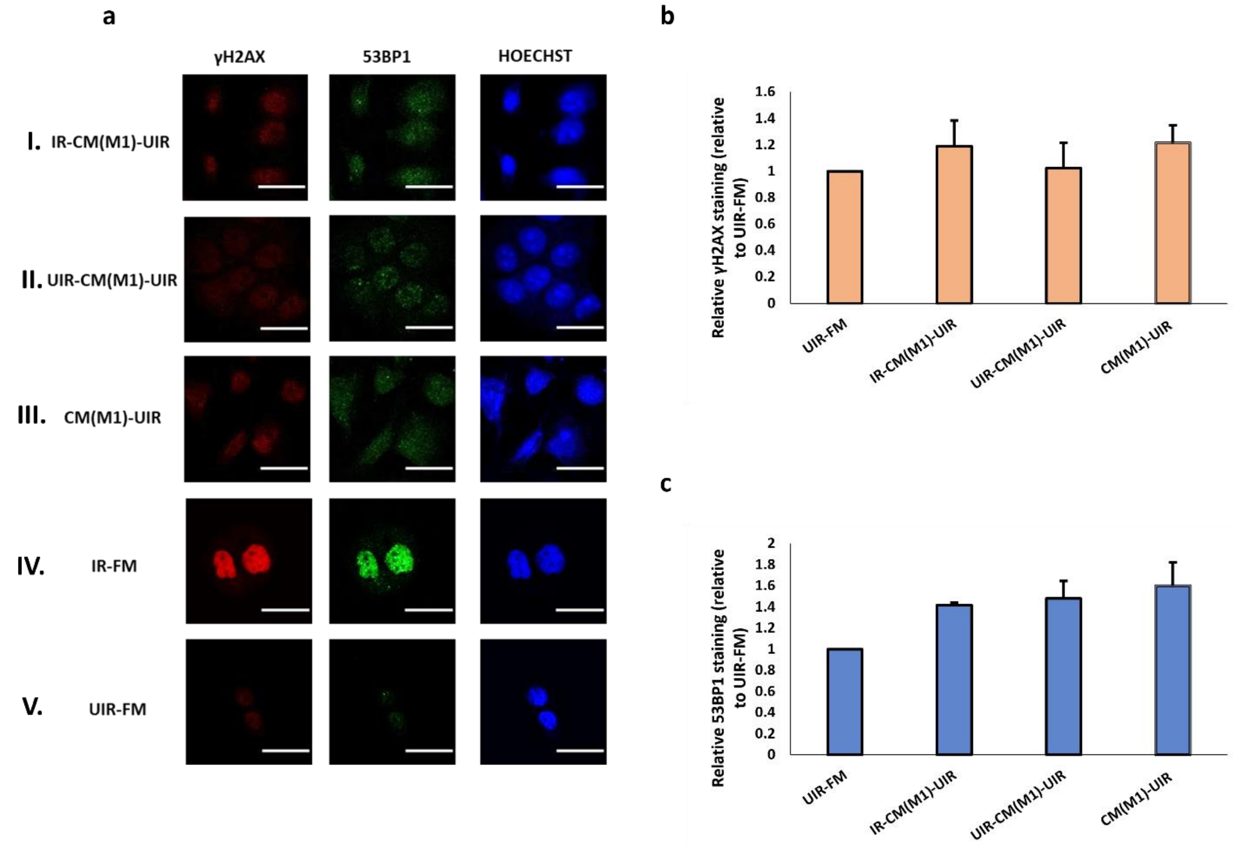


**Supplementary Fig. 4** (a) Representative images from immunofluorescence staining with anti-phospho-histone H2A.X and anti-53BP1 antibodies in MCF7 cells post-UVC mock-irradiation. CM experiments with M1 cells as Bystanders were carried out in the following conditions, (I) IR-CM(M1)-UIR, (II) UIR-CM(M1)-UIR, (III) CM(M1)-UIR. Control experiments included (IV) IR-FM and (V) UIR-FM. Scale bar = 25 μm. (b) Graph representing relative γH2AX staining (linear values relative to UIR-FM) for these conditions. (c) Graph representing relative 53BP1 staining (linear values relative to UIR-FM) for these conditions. * $P < 0.05$; ** $P < 0.01$ and error bars represent mean ± SD.

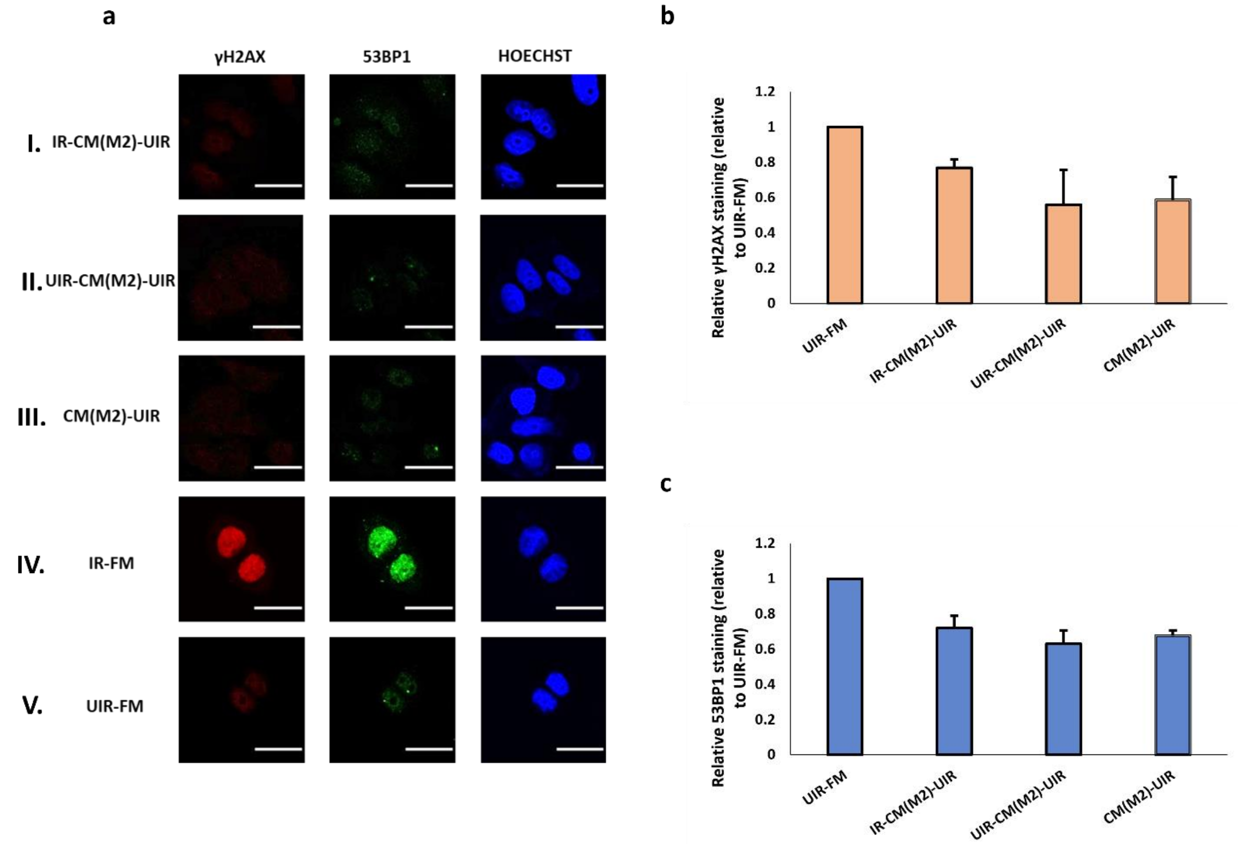


**Supplementary Fig. 5** (a) Representative images from immunofluorescence staining with anti-phospho-histone H2A.X and anti-53BP1 antibodies in MCF7 cells post-UVC mock-irradiation. CM experiments with M2 cells as Bystanders were carried out in the following conditions, (I) IR-CM(M2)-UIR, (II) UIR-CM(M2)-UIR, (III) CM(M2)-UIR. Control experiments included (IV) IR-FM and (V) UIR-FM. Scale bar = 25 μm. (b) Graph representing relative γH2AX staining (linear values relative to UIR-FM) for these conditions. (c) Graph representing relative 53BP1 staining (linear values relative to UIR-FM) for these conditions. * $P < 0.05$; ** $P < 0.01$ and error bars represent mean ± SD.

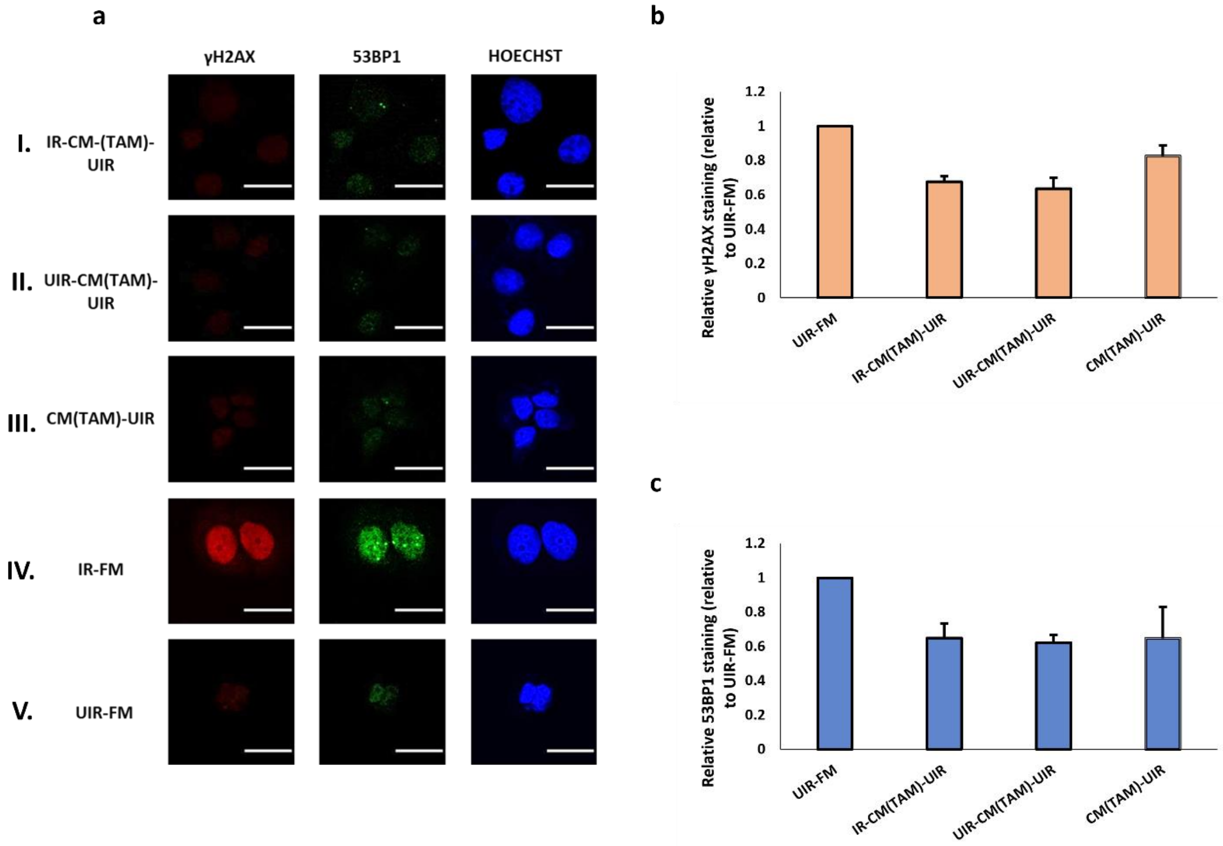


**Supplementary Fig. 6** (A) Representative images from immunofluorescence staining with anti-phospho-histone H2A.X and anti-53BP1 antibodies in MCF7 cells post-UVC mock-irradiation. CM experiments with TAMs as Bystanders were carried out in the following conditions, (I) IR-CM(TAM)-UIR, (II) UIR-CM(TAM)-UIR, (III) CM(TAM)-UIR. Control experiments included (IV) IR-FM and (V) UIR-FM. Scale bar = 25 µm. (B) Graph representing relative γH2AX staining (linear values relative to UIR-FM) for these conditions. (C) Graph representing relative 53BP1 staining (linear values relative to UIR-FM) for these conditions. * $P < 0.05$; ** $P < 0.01$ and error bars represent mean ± SD
